# An extension of the Walsh-Hadamard transform to calculate and model epistasis in genetic landscapes of arbitrary shape and complexity

**DOI:** 10.1101/2023.03.06.531391

**Authors:** Andre J. Faure, Ben Lehner, Verónica Miró Pina, Claudia Serrano Colome, Donate Weghorn

## Abstract

Accurate models describing the relationship between genotype and phenotype are necessary in order to understand and predict how mutations to biological sequences affect the fitness and evolution of living organisms. The apparent abundance of epistasis (genetic interactions), both between and within genes, complicates this task and how to build mechanistic models that incorporate epistatic coefficients (genetic interaction terms) is an open question. The Walsh-Hadamard transform represents a rigorous computational framework for calculating and modeling epistatic interactions at the level of individual genotypic values (known as genetical, biological or physiological epistasis), and can therefore be used to address fundamental questions related to sequence-to-function encodings. However, one of its main limitations is that it can only accommodate two alleles (amino acid or nucleotide states) per sequence position. In this paper we provide an extension of the Walsh-Hadamard transform that allows the calculation and modeling of background-averaged epistasis (also known as ensemble epistasis) in genetic landscapes with an arbitrary number of states per position (20 for amino acids, 4 for nucleotides, etc.). We also provide a recursive formula for the inverse matrix and then derive formulae to directly extract any element of either matrix without having to rely on the computationally intensive task of constructing or inverting large matrices. Finally, we demonstrate the utility of our theory by using it to model epistasis within both simulated and empirical multiallelic fitness landscapes, revealing that both pairwise and higher-order genetic interactions are enriched between physically interacting positions.

**Author Summary:** An important question in genetics is how the effects of mutations combine to alter phenotypes. Genetic interactions (epistasis) describe non-additive effects of pairs of mutations, but can also involve higher-order (three- and four-way etc.) combinations. Quantifying higher-order interactions is experimentally very challenging requiring a large number of measurements. Techniques based on deep mutational scanning (DMS) represent valuable sources of data to study epistasis. However, the best way to extract the relevant pairwise and higher-order epistatic coefficients (genetic interaction terms) from this data for the task of phenotypic prediction remains an unresolved problem. The Walsh-Hadamard transform represents a rigorous computational framework for calculating and modeling epistatic interactions at the level of individual genotypic values. Critically, this formalism currently only allows for two alleles (amino acid or nucleotide states) per sequence position, hampering applications in more biologically realistic scenarios. Here we present an extension of the Walsh-Hadamard transform that overcomes this limitation and demonstrate the utility of our theory by using it to model epistasis within both simulated and empirical multiallelic genetic landscapes.

## Introduction

A fundamental challenge in biology is to understand and predict how changes (or mutations) to biological sequences (DNA, RNA, proteins) affect their molecular function and ultimately the phenotype of living organisms. The phenomenon of ‘epistasis’ (genetic interactions) – broadly defined as the dependence of mutational effects on the genetic context in which they occur [1, 2, 3] – is widespread in biological systems, yet knowledge of the underlying mechanisms remains limited. Defining the extent of epistasis and better understanding of its origins has relevance in fields ranging from genetic prediction, molecular evolution, infectious disease and cancer drug development, to biomolecular structure determination and protein engineering [3].

Evolutionarily related sequences, natural genetic variation within populations, and more recently results of techniques such as deep mutational scanning (DMS) [4] – also known as massively parallel reporter assays (MPRAs) and multiplex assays of variant effect (MAVEs) – represent valuable sources of data to study epistasis [5, 1]. In particular, DMS enables the systematic measurement of mutational effects across entire combinatorially complete genetic landscapes [5, 6, 7, 8, 9, 10, 11, 12, 13]. Importantly, the typical use of engineered genotypes, haploid individuals and near-identical environmental (laboratory) conditions in these experiments allows population genetic considerations – such as dominance, variable allele frequencies and linkage disequilibrium – to be ignored [14]. In other words, measurements obtained from deep mutational scanning and related methods permit the modeling of epistasis in the mechanistic sense (sequence-to-function encoding) rather than in the evolutionary sense, i.e. based on the dynamics of population genotype frequencies. Nevertheless, precisely how to extract the most biologically relevant pairwise and higher-order epistatic coefficients (genetic interaction terms) from this type of data is an unresolved problem.

Quantitative definitions of epistasis vary among fields, but it has been argued that one particular formulation termed ‘background-averaged’ epistasis, also known as ‘ensemble’ epistasis [1, 12], may provide the most useful information on the epistatic structure of biological systems [2]. The underlying rationale is that by averaging the effects of mutations across many different genetic backgrounds (contexts), the method is robust to local idiosyncrasies in the relationship between genotype and phenotype. It has been previously pointed out that the definition of background-averaged epistasis is conceptually similar to that of ‘statistical epistasis’ attributed to Fisher, but instead of measuring the average effect of allele substitutions against the population average genetic background i.e. averaging over all genotypes present in a given population (taking into account their individual frequencies), the approach instead averages over all possible genotypes (assuming equal genotype weights) [1, 2].

The current mathematical formalism of background-averaged epistasis is based on the Walsh-Hadamard transform [2]. Interestingly, although widely used in physics and engineering, the Walsh-Hadamard transform was first applied to non-biological fitness landscapes in the field of genetic algorithms (GA) [15], subsequently being proposed as the basis of a framework for the computation of higher-order epistasis in empirical settings [16]. However, the Walsh-Hadamard transform can only accommodate two alleles (amino acid or nucleotide states) per sequence position, with no extension to multialleleic landscapes (cardinality greater than two) yet made, as confirmed by multiple recent reports [2, 17, 18, 19]. Alternative implementations for multiallelic landscapes either rely on ‘one-hot encoding’ elements of larger alphabets as biallelic sequences – requiring the manipulation of prohibitively large Walsh-Hadamard matrices – or constructing graph Fourier bases [18, 20], which is mathematically complex and provides no straightforward way to interpret epistatic coefficients. The result is that the application of background-averaged epistasis has been severely limited and its properties remain largely unexplored in more biologically realistic scenarios.

In this work we provide an extension of the Walsh-Hadamard transform that allows the calculation and modeling of background-averaged epistasis in genetic landscapes with an arbitrary number of states (20 for amino acids, 4 for nucleotides, etc.). We also provide a recursive formula for the inverse matrix, which is required to infer epistatic coefficients using regression. Furthermore, we derive convenient formulae to directly extract any element of either matrix without having to rely on the computationally intensive task of constructing or inverting large matrices. Lastly, we apply these formulae to the analysis of both simulated and empirical multiallelic DMS datasets, demonstrating that sparse models inferred from the background-averaged representation (embedding) of the underlying genetic landscape more regularly include epistatic terms corresponding to direct physical interactions.

## Results

### Extension of the Walsh-Hadamard transform to multiallelic landscapes

In this work, a genotype sequence is represented as a one-dimensional ordering of monomers, each of which can take on *s* possible states (or alleles), for example *s* = 4 for nucleotide sequences or *s* = 20 for amino acid sequences. Without loss of generality, the *s* states can be labelled 0, 1, 2, …, *s* − 1, where 0 denotes the wild-type allele. We are going to consider genotype sequences of length *n* ∈ ℕ, i.e. sequences taking values in 𝒮^*n*^, where 𝒮:= {0, 1, …, *s* − 1}.

Each genotype 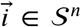 is associated with its phenotype 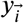. Note that here we use the term ‘phenotype’ as shorthand for ‘molecular phenotype score’ from a quantitative laboratory assay (DMS) reporting on a molecular function for each genotype of interest. In quantitative genetics terminology this might be referred to as ‘genotypic value’ because environmental deviation is negligible due to the controlled nature of the experiments, but our subject here is the macromolecule not an individual from a population [14]. In the context of empirical genotype-phenotype landscapes, the phenotypic effect of a genotype 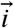 is typically measured with respect to the wild-type, i.e. it is given by 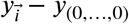.

It is important to emphasize that in what follows we implicitly restrict ourselves to the haploid reference base, because our primary goal is the modeling of sequence-to-function encodings for *individual* genotype sequences – for the ultimate purpose of understanding and engineering macromolecules – not the modeling of sequence evolution or quantification of sources of phenotypic variance in populations.

If the phenotypic effects of individual mutations were independent, they would be additive, meaning that the phenotypic effect of 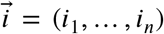 would be the sum of the phenotypic effects of the single mutants (*i*_1_, 0, …, 0), …, (0, …, 0, *i*_*n*_). The epistatic coefficient quantifies how much the observed phenotypic effect of 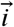 deviates from this assumption. In the case of background-averaged epistasis, we quantify the interactions between a set of mutations by averaging over all possible genotypes for the remaining positions in the sequence. For example, if *n* = 3 and *s* = 2, the pairwise epistatic coefficient involving the mutations at positions 2 and 3 is calculated by averaging over all states (backgrounds) for the remaining positions, in this case given by the two states of the first position (* denotes the positions at which the averaging is performed), i.e.

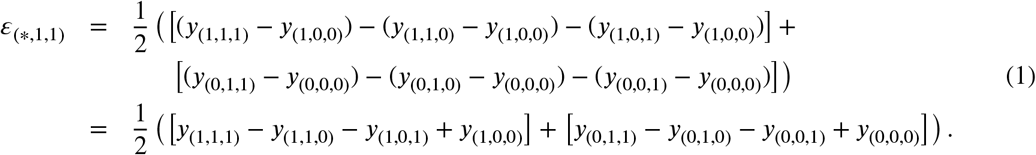

More generally, in [2] it is shown that for *s* = 2 and any sequence length *n*, phenotypic effects can be decomposed into background-averaged epistatic coefficients with

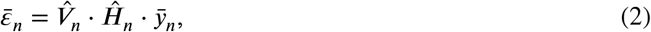

where 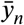 is the vector 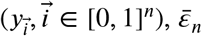 is the vector 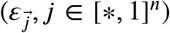 and Ĥ_*n*_ and 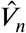 are 2^*n*^ × 2^*n*^matrices defined recursively as follows:

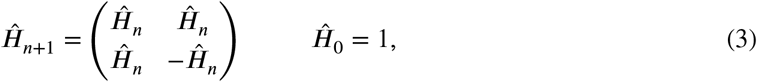

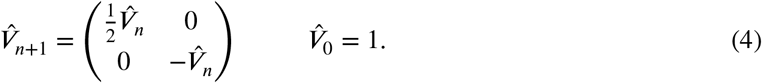

The matrix Ĥ is known as the Walsh-Hadamard transform [21, 22] and 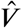 is a diagonal weighting (or normalisation) matrix to correct the sign and account for averaging over different numbers of backgrounds as a function of epistatic order [2].

In this work, we provide an extension of this theory to describe background-averaged epistasis for sequences with an arbitrary number of states *s*. Before writing a general formula, we consider the simplest possible multi-state (multiallelic) landscape i.e. a sequence of length *n* = 1 with *s* = 3,

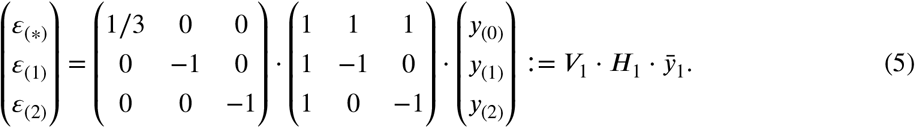

Consistent with the definition of background-averaged epistasis for biallelic landscapes [2], the zeroth-order epistatic coefficient ε_(*)_ is the mean phenotypic value across all genotypes and the first-order epistatic coefficients ε_(1)_ and ε_(2)_ are simply the respective individual phenotypic effects of genotypes *y*_(1)_ and *y*_(2)_ with respect to the wild-type. However, the key feature of *H*_1_ for multiallelic landscapes – and where it departs from the canonical Walsh-Hadamard transform – is the introduction of zero elements to exclude phenotypes that are irrelevant for the calculation of a given epistatic coefficient. In other words, these phenotypes are excluded because they correspond neither to relevant intermediate genotypes nor alternative genetic backgrounds.

If we now consider a sequence of length *n* = 2 with *s* = 3, then the *H*_2_ and *V*_2_ matrices become 9 ×9 (*s*^*n*^×*s*^*n*^) and can be constructed from recurring to the case *n* = 1 above, giving

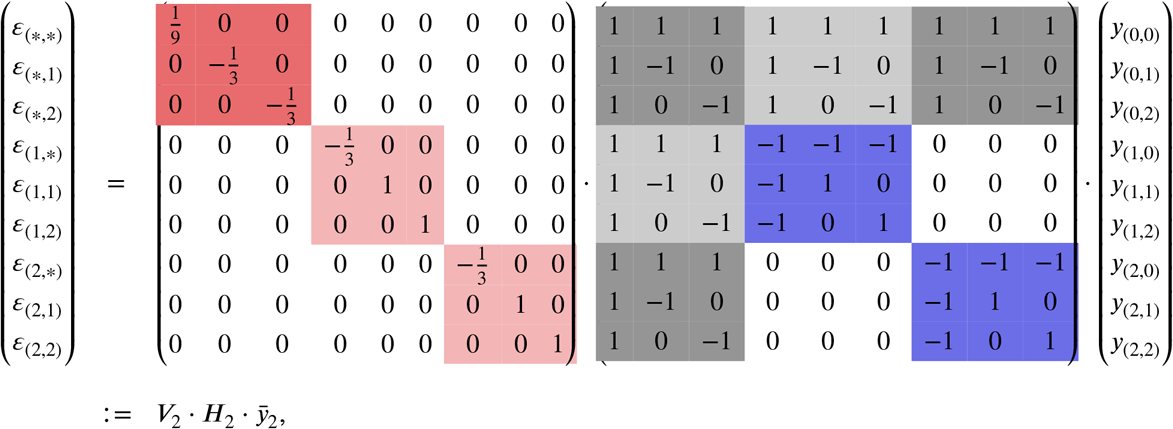

where the colors highlight the block structure of the matrices. In *V*_2_, the red square corresponds to 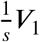and the light red squares to −*V*_1_. In *H*_2_, the gray squares correspond to *H*_1_ and the blue squares to −*H*_1_. In Table 1 we show the results of background-averaged epistatic coefficients calculated by applying the above formula to an empirical multiallelic landscape with *n* = 2 and *s* = 3 [6].

**Table 1:** Interaction terms based on background-averaged epistasis 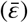 for an empirical multiallelic genotype-phenotype landscape consisting of all combinations of two mutations each at positions 6 and 66 in the tRNA-Arg(CCU) [6], i.e. *n* = 2 and s= 3. The first two columns indicate nucleic acid sequences and their base 3 representations. Here the ‘GC’ reference (wild-type) genotype corresponds to that of *S. cerevisiae*, denoted by (0 0). The second two columns show the measured phenotypic effects and corresponding background-averaged epistatic coefficients together with their empirical errors and their propagated errors. See Results for a regression analysis of the entire dataset and Supplementary Methods for a derivation of the error propagation.

| Nucleic acid sequence | Base $s = 3$ representation | Phenotypic effect $\bar{y}_2$ | Epistatic term $\bar{\epsilon}_2 = V_2 \cdot H_2 \cdot \bar{y}_2$ |
| --- | --- | --- | --- |
| GC | (0,0) | 0 ($\pm 0.01$) | -0.17 ($\pm 0.02$) |
| GA | (0,1) | -0.14 ($\pm 0.05$) | -0.21 ($\pm 0.06$) |
| GT | (0,2) | -0.07 ($\pm 0.02$) | 0.02 ($\pm 0.02$) |
| AC | (1,0) | -0.13 ($\pm 0.02$) | -0.24 ($\pm 0.05$) |
| AA | (1,1) | -0.8 ($\pm 0.15$) | -0.53 ($\pm 0.16$) |
| AT | (1,2) | -0.01 ($\pm 0.03$) | 0.19 ($\pm 0.04$) |
| TC | (2,0) | -0.19 ($\pm 0.02$) | -0.05 ($\pm 0.03$) |
| TA | (2,1) | 0 ($\pm 0.04$) | 0.33 ($\pm 0.07$) |
| TT | (2,2) | -0.18 ($\pm 0.03$) | 0.08 ($\pm 0.04$) |

More generally, for any value of *s*, when *n* = 1,

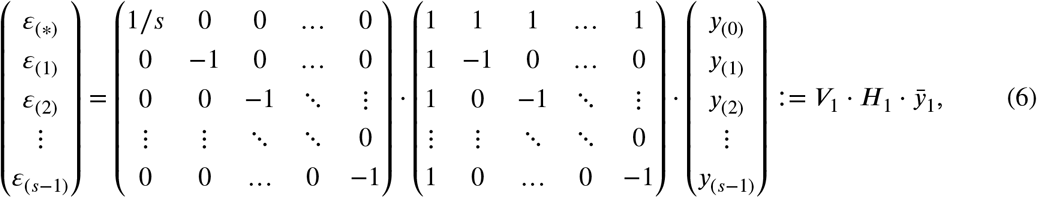

where ε_(*)_ corresponds to averaging phenotypes over all possible genotypes and the remaining coefficients simply correspond to the phenotypic effects of each mutation.

For *n* = 2, we have to consider different combinations of mutations in both positions. In this case, the phenotypes can be written as

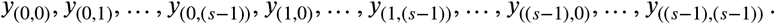

A natural ordering of the phenotypes is given by interpreting genotype 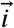 as the base *s* representation of an integer (see Table 1). From this, we can see how the first *s* genotypes correspond to combining the wild-type allele at the first position with a state from the case *n* = 1, i.e. to genotypes that can be written 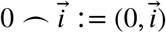, with 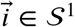. The next *s* genotypes correspond to the first mutated allele at the first position combined with all the genotypes of *n* = 1, i.e. 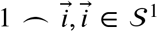, and so on. Therefore, we can write the matrices *H* and *V* following a block structure. In the case *n* = 2 and any given *s*, we would then have

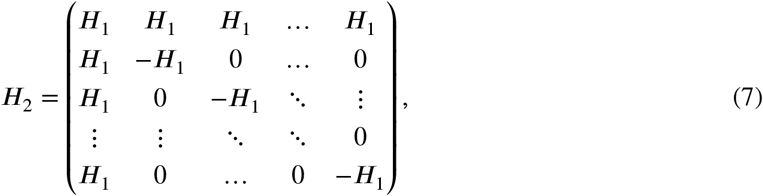

where the number of *H*_1_ blocks corresponds to the number of states of the first position, so *s*. Moreover, each of these blocks must be normalized to yield the corresponding background-averaged epistatic terms. Therefore *V*_2_ can also be expressed as a function of *V*_1_ as follows:

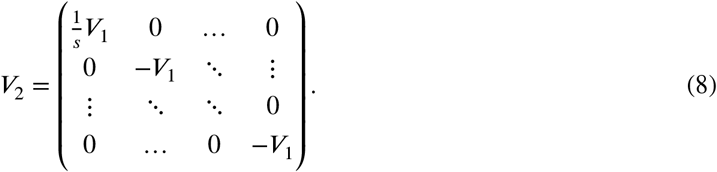

Given these two matrices, we can write the background-averaged epistatic coefficients for the case of *n* = 2 and *s* different states per position as 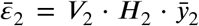. More generally, the decomposition of phenotypic effects into background-averaged epistatic coefficients is given by

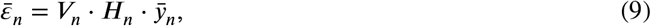

where *H*_*n*_ and *V*_*n*_ can be defined recursively as

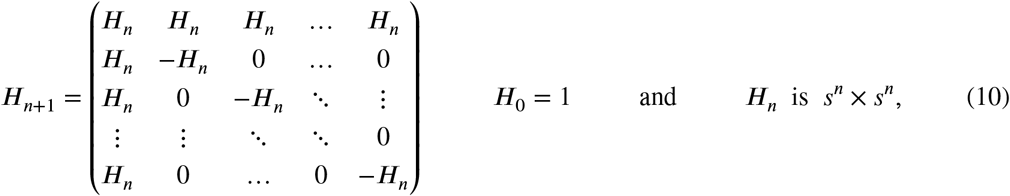

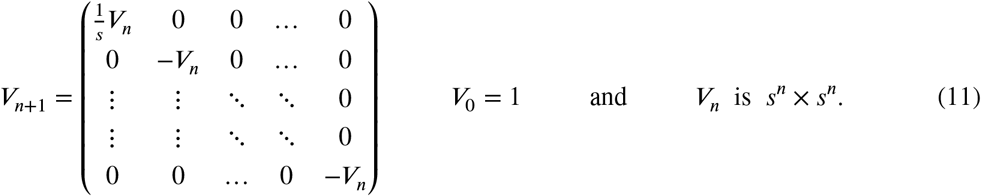

### Recursive inverse matrix

Equation (9) defines the vector of epistatic coefficients, 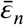, as a function of the vector of phenotypes, 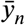, which in general is the quantity that is measured experimentally. However, usually phenotypic measurements are only available for a subset of genotypes. An alternative is therefore to estimate the epistatic coefficients 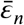 by regression,

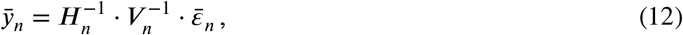

where the product 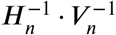 represents a matrix of sequence features. This is analogous to the more widely used one-hot encoding strategy, which implicitly relies on a ‘background-relative’ (or ‘biochemical’) view of epistasis when regressing to full order [2]. We discuss other advantages of estimating background-averaged epistatic coefficients using regression at the end of this manuscript.

Since *V*_*n*_ is a diagonal matrix, its inverse is also a diagonal matrix whose elements are the inverse of the elements of *V*_*n*_.

The inverse of *H*_*n*_ is the matrix *A*_*n*_ which can be defined recursively as

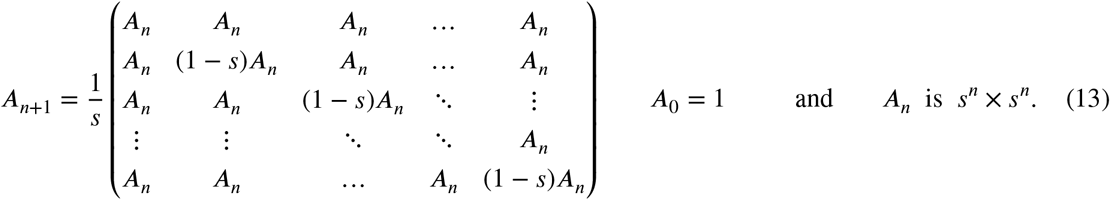

See Proposition 1 in S1 Text for a proof of this result. This is the most efficient method to determine the full matrix *A*_*n*_ (see Results).

### Formulae to obtain elements of the matrices

When regressing phenotypes on genotypes, a common goal is to determine whether epistatic coefficients up to the *r*^*th*^ order (where *r* < *n*) are sufficient to describe the complexity of the biological system. Furthermore, as mentioned above, some fraction of phenotype values within combinatorially complete genetic landscapes are typically unavailable, representing missing data. Restricting the epistatic order and missing phenotypes respectively correspond to omitting rows and columns from *H*_*n*_ (and vice versa from *A*_*n*_). Formulae to directly obtain elements of the matrices in equations (9) and (12) would therefore be convenient.

In order to write the matrix element (*H*_*n*_)_*ij*_, we need to compare the genotype sequences,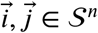,

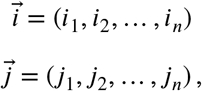

where 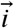 denotes the *i*^*th*^element in 𝒮^*n*^, 𝒮= {0, 1, …, *s* − 1}, and the elements of 𝒮^*n*^ are ordered by the base *s* representation of integers. For instance, for any value of *n*, we will denote the wild-type state with index *i* = 1 and write 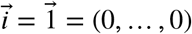. The element denoted with index *i* = 2 would be 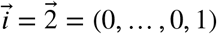 and so on.

The elements of *H*_*n*_ can be written as

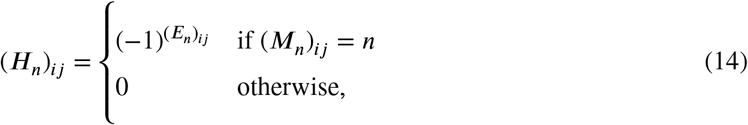

where *M* and *E* are *s*^*n*^× *s*^*n*^matrices whose elements are

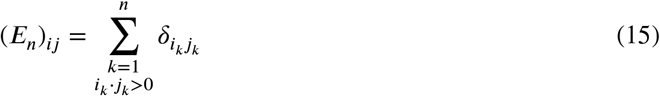

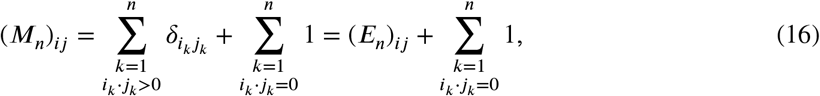

where δ_*ij*_ denotes the Kronecker delta of *i, j*, which is equal to 1 when *i* = *j* and 0 if *i* ≠ *j*. In words, (*E*_*n*_)_*ij*_ counts the number of positions at which the genotype sequences 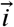 and 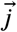 carry the same mutated allele and (*M*_*n*_)_*ij*_ is equal to (*E*_*n*_)_*ij*_ plus the number of positions where 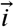 or 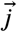 carry the wild-type allele. See Proposition 2 in S1 Text for a proof of this result.

Furthermore, the elements of *A*_*n*_ can be written as

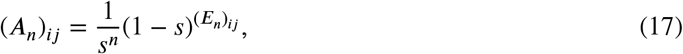

where *E*_*n*_ is defined as in (15). See Proposition 3 in S1 Text for a proof of this result.

Finally, the matrices *V*_*n*_ and 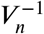 are diagonal matrices whose diagonal elements can be written as

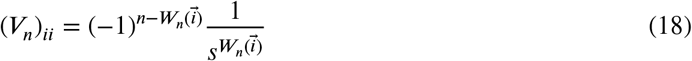

and

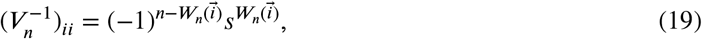

where

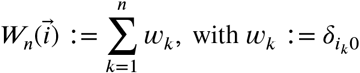

and 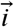 again denotes the *i*^*th*^ element in 𝒮^*n*^ when ordered by the base *s* representation of integers. In words, *w*_*k*_ = 1 if the genotype sequence 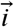 carries the wild-type allele at position *k* and 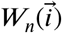 counts the number of positions in 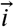 carrying the wild-type allele. We prove this result in Proposition 4 in S1 Text.

### Generalization to different numbers of states per position

We can generalize the formulae described in the previous subsection further by considering that each position can have different numbers of states. In this case, we can denote *s*_*k*_ the number of possible states at position *k*. For *n* = 1, this corresponds to exactly the same matrix as in the previous case but with *s* = *s*_1_, which is the number of possible states in this position. For *n* = 2, the matrix changes because now the new position can have a different number of possible states, *s*_2_. Following the recursive definition of *H*_*n*_, we can construct *H*_2_ by repeating *H*_1_ *s*_2_ times, with the structure stated in (10). Therefore, we have

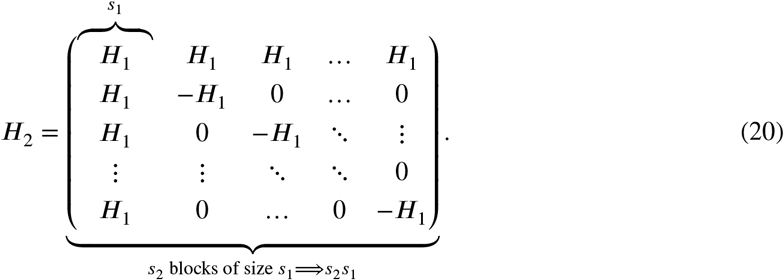

So the structure is exactly the same but the size of the matrix for each *n* varies according to the number of possible states of the new position. The definition of *H*_*n*_ is the same as in (10) but the dimensions of the matrix are 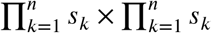. Similarly, the inverse matrix *A*_*n*+1_ can be written recursively as

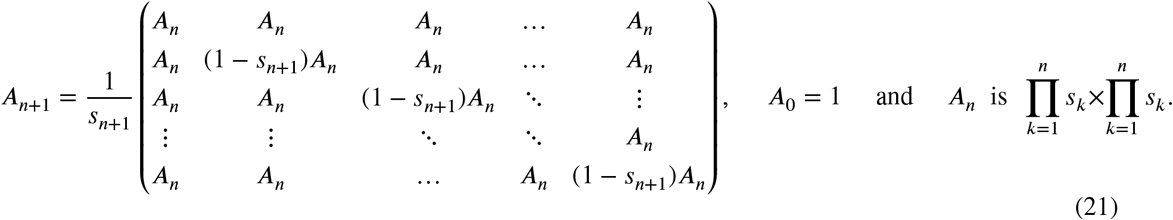

The matrix *A*_*n*_ defined in (21) is the inverse of the matrix *H*_*n*_ in the general case where each position can have a different number of states.

In this general case, the elements of *H*_*n*_ and *A*_*n*_ can be written as

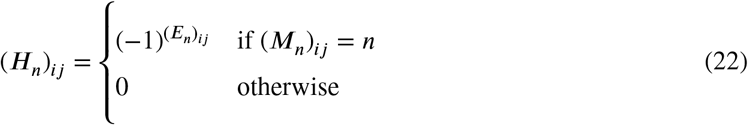

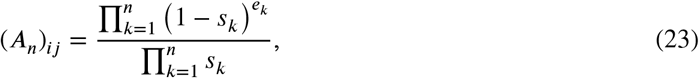

where *E*_*n*_ and *M*_*n*_ are defined as in (15) and 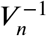.

The matrices *V*_*n*_ and 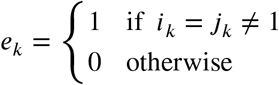 are diagonal matrices whose diagonal elements can be written as

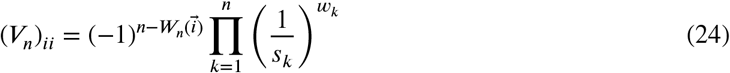

and

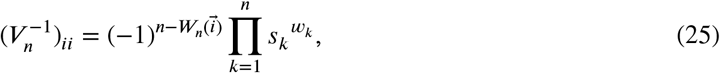

where

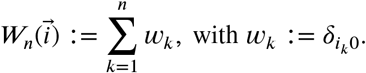

We prove the results in this subsection in Propositions 5, 6 and 7 in S1 Text.

The above formulae permit the calculation and modeling of background-averaged epistasis in arbitrarily-shaped genetic landscapes, i.e. with any number of alleles (states) per position, as well as the direct construction of sub-matrices for regression to any desired epistatic order and/or in the presence of missing data. In the following subsections we report benchmarking results comparing the performance of alternative methods to obtain *H*_*n*_ and *A*_*n*_, as well as results from the application of our theory extension to an empirical multiallelic genotype-phenotype landscape.

### Benchmarking

Fig 1a-d provides a visualization of the matrices *H*_*n*_ and *A*_*n*_ for different values of *n* and *s*, clearly showing a self-similar pattern in all cases due to their recursive nature.

**Figure 1:**
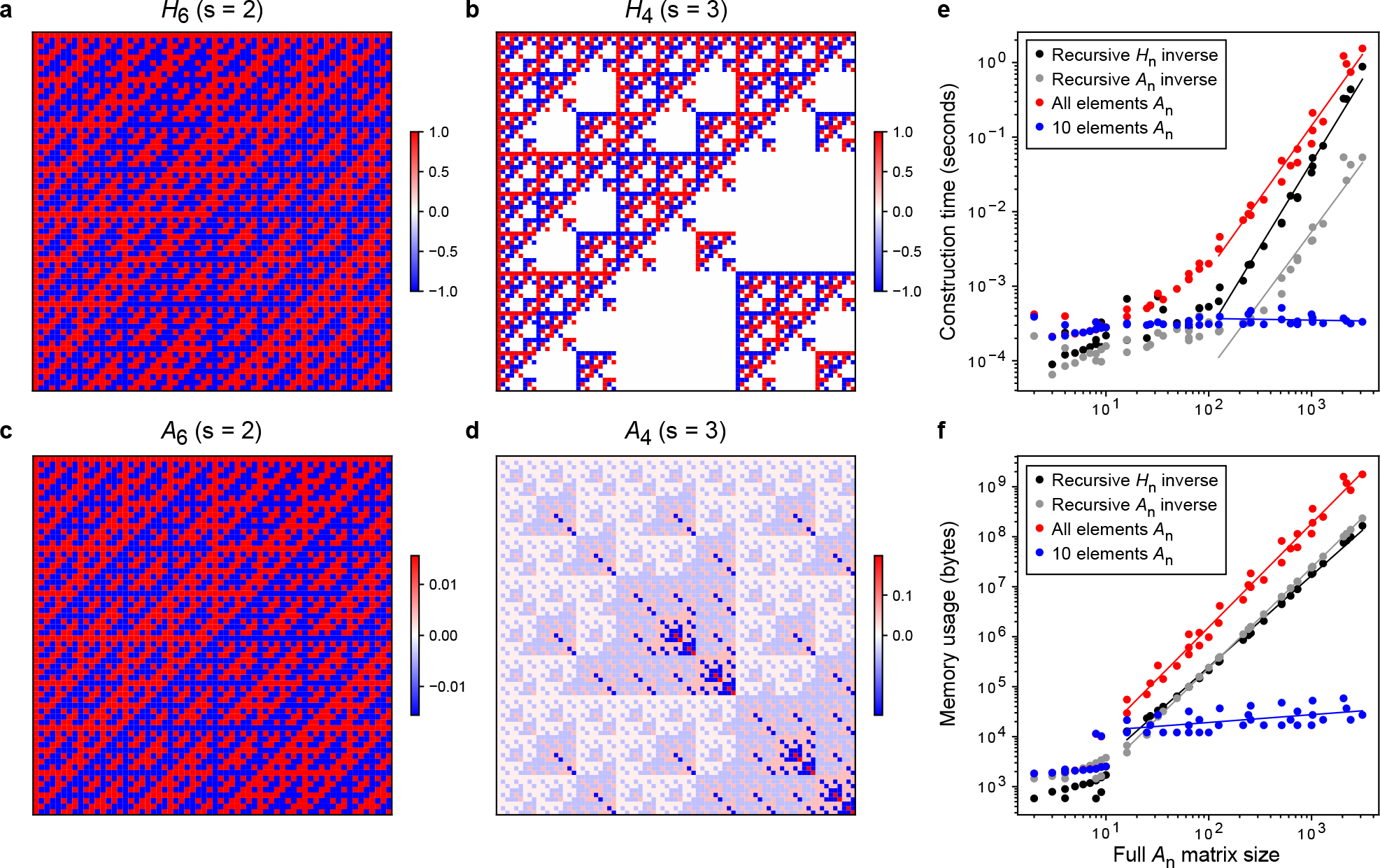
Benchmarking results and heat map representations of matrices corresponding to the binary (biallelic) and multi-state (multiallelic) extension of the Walsh-Hadamard transform, and their corresponding inverses. **a**, *H*_6_ Walsh-Hadamard transform (s=2). **b**, *H*_4_ multi-state extension of the Walsh-Hadamard transform for *s* = 3. **c**, *A*_6_ Inverse Walsh-Hadamard transform. **d**, *A*_4_ multi-state extension of the inverse Walsh-Hadamard transform for *s* = 3. **e**, Computational time on a MacBook Pro (13-inch, 2017, 2.3GHz dual-core Intel Core i5) for extracting elements of *A*_*n*_ matrices of various dimensions and numbers of states (alleles) per position (*s* ∈ [2, 10]). Comparisons are shown between numerically inverting the recursively constructed *H*_*n*_ (using scipy.linalg.inv), i.e. “Recursive *H*_*n*_ inverse”, using the recursive formula for *A*_*n*_, using the formula to extract all elements of *A*_*n*_ and extracting 10 random elements of *A*_*n*_ (see legend). The mean across 10 replicates is depicted. Linear regression lines were fit to data from matrices with at least 100 elements. **f**, Similar to **e** but indicating memory usage. Linear regression lines were fit to data from matrices with at least 10 elements.

In this paper, we provide different methods to construct 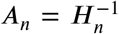. First, *H*_*n*_ can be numerically inverted using standard matrix inversion algorithms (here we use the linalg.inv function from the SciPy library in Python), referred to as “Recursive *H*_*n*_ inverse” in Fig 1e,f. Alternatively, the recursive definition of the inverse given by equation (13) can be used, which we refer to as “Recursive *A*_*n*_”. As can be seen in Fig 1e, this method is faster than numerically inverting *H*_*n*_.

Finally, we also provide a convenient formula for extracting specific individual elements of *A*_*n*_ (Proposition 3), referred to as “All elements *A*_*n*_” in Fig 1e,f. This method is more computationally intensive than the previously described methods, due to the formula relying on the computation of (*E*_*n*_)_*ij*_, which equates to counting the number of sequence positions that are identically mutated in vectors 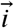 and 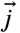, each of size *n*. However, in situations where subsets of elements (or sub-matrices) – rather than full matrices – are desired, Proposition 3 provides a method that can be faster and more memory efficient (see “10 elements *A*_*n*_” in Fig 1e,f).

For example, in the case of a 10-mer DNA sequence, constructing the full inverse transform *A*_10_ with *s* = 4 would require > 10^23^ bytes (100 million petabytes) of memory in the best-case scenario (“Recursive *H*_*n*_ inverse” in Fig 1f, log-linear extrapolation). Similarly, the full inverse transform for a 4-mer amino acid sequence (*A*_4_ with *s* = 20) would impose a memory footprint > 10^20^ bytes. On the other hand, calculating the subset of elements from these matrices required for the prediction of a single phenotype using epistatic coefficients up to third order (three-way genetic interaction terms) is feasible in both situations using Proposition 3 (3,675 and 29,678 elements; 2.5 GB and 192 GB of memory; 1.8 and 99 seconds, respectively). This memory footprint can easily be diminished further using data chunking, which is a unique benefit of this method.

### Application to a simulated multiallelic genotype-phenotype landscape

In order to demonstrate the utility of our extension of the Walsh-Hadamard transform, we used it to model epistasis within a simulated multiallelic genetic landscape. Knowledge of the ‘ground truth’ epistatic coefficients allowed us to (i) determine whether background-averaged coefficients can be accurately inferred using regression and (ii) investigate the impact of data sparseness on the results i.e. in the context of missing phenotypic values. In most practical applications – particularly in the case of large empirical genotype-phenotype landscapes – measuring the phenotypic effects of all mutation combinations is infeasible. The simulated combinatorially complete landscape consisted of all possible combinations of four alleles at six different positions i.e. a total of 4^6^ = 4, 098 genotypes (Fig 2a, left). First-order and a subset of second-order epistatic coefficients were randomly drawn from Gaussian distributions, while all higher-order epistatic coefficients were set to zero (see Methods). We arbitrarily selected three pairs of positions at which mutations genetically interact i.e. ground truth non-zero pairwise epistatic terms (Fig 2a, left). Fitness values for all variants were reconstructed using equation (12) with random noise added to simulate measurement error (see Methods).

**Figure 2:**
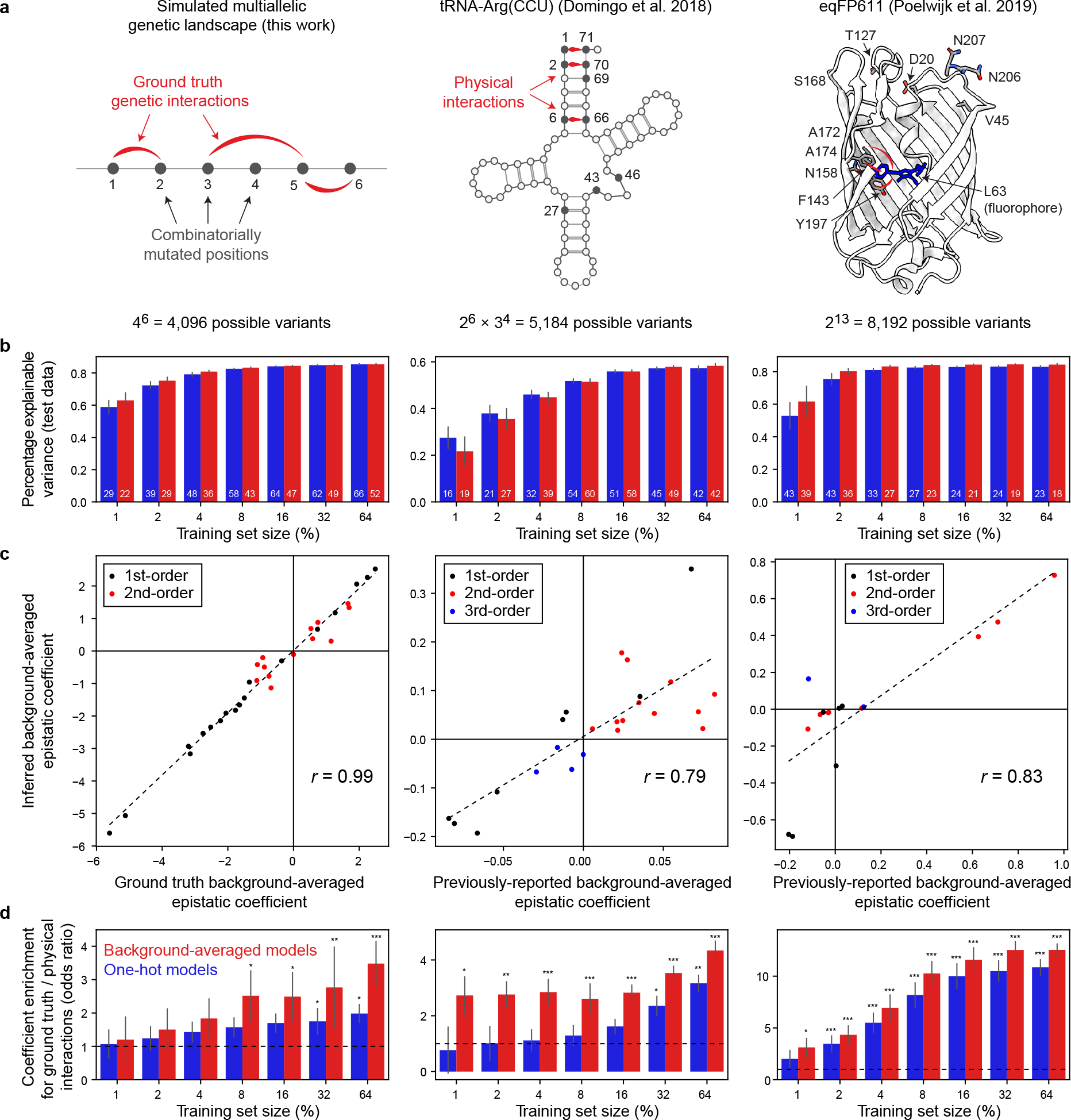
Learning sparse models from simulated and empirical combinatorial fitness landscapes. **a**, Schematic description of combinatorial DMS datasets. Left: simulated multiallelic landscape consisting of all combinations of 4 nucleotides in a DNA 6-mer where ground truth interactions ([1,2], [3,5] and [5,6]) are depicted with red arcs (see Methods). Middle: secondary structure of *S. cerevisiae* tRNA-Arg(CCU) indicating variable positions (closed circles) combinatorially mutated in a DMS experiment as described in [6]. Three Watson–Crick base pairing (WCBP) interactions involving pairs of these positions ([1,71], [2,70] and [6,66]) are indicated. Right: crystal structure of a blue fluorescent variant (TagBFP) of the *Entacmaea quadricolor* protein eqFP611 (PDB: 3M24) and 13 positions (12 shown) that differ in the red fluorescent variant (mKate2) that were mutated in a DMS experiment as described in [7]. Physical interactions between the fluorophore (L63) and three proximal residues ([63,143], [63,174] and [63,197]) are indicated. **b**, Performance of sparse models fit using different proportions of the DMS datasets indicated in panel a. The median number of model coefficients is indicated. Colour scale as in panel d. **c**, Scatter plots comparing sparse model-inferred background-averaged epistatic coefficients (training set size = 64%) to ground truth values (left panel) or previously-reported coefficients (remaining panels) [6, 7]. **d**, Enrichment of direct physical interactions (red arcs in panel a) in non-zero epistatic coefficients. *, *P* < 0.05; **, *P* < 0.01; ***, *P* < 0.001. Error bars indicate the interquartile range.

We then trained Lasso regression models of the form in equation (12) to predict fitness values from sequence, where the inferred model parameters correspond to background-averaged epistatic coefficients up to third order (S1a Fig; see Methods). To assess the impact of data sparsity on the results, we sub-sampled the simulated phenotype (fitness) values to obtain training dataset sizes ranging from 64% to 1% of all variants in the combinatorial complete landscape. The sparse models generalize well, explaining more than 80% of fitness variance in the held out test data even when the overwhelming majority of variants (96%) are missing (Fig 2b, left; ‘Background-averaged models’, red). Importantly, the values of non-zero background-averaged epistatic coefficients closely match those of the ground truth simulated coefficients (Pearson’s *r* = 0.99, Fig 2c, left). Reducing the size of the training set tends to reduce the number of coefficients retained in sparse models, but the ground truth values of non-zero coefficients are well recovered (Pearson’s *r* = 0.99, S2a Fig).

For comparison, we used the same procedure to fit Lasso regression models of the form 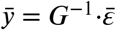, where *G*^−1^ represents a matrix of one-hot encoded sequence features i.e. the presence or absence of a given mutation – or mutation combination (interaction) – with respect to the reference (wild-type) genotype is denoted by a ‘1’ or ‘0’ respectively (S1a Fig, Fig 2b, left; ‘One-hot models’, blue). The definition of *G* and its relationship to the biochemical (or background-relative) view of epistasis is explained in [2]. Although one-hot models show similar performance to background-averaged models on simulated test data, they tend to be more complex with larger numbers of coefficients (Fig 2b), including spurious third-order epistatic terms although their ground truth values are zero (S1c Fig).

In summary, these results demonstrate that our theory extensions allow the accurate inference of background-averaged epistatic coefficients in multiallelic genetic landscapes, even in situations when large amounts of phenotypic data are missing.

### Application to empirical combinatorial fitness landscapes

We next applied our theory to model epistasis within two different empirical combinatorial fitness landscapes (Fig 2a, middle, right). In the first DMS assay, a budding yeast strain was used in which the single-copy arginine-CCU tRNA (tRNA-Arg(CCU)) gene is conditionally required for growth [6]. A library of variants of this gene was designed to cover all 5,184 (2^6^ ×3^4^) combinations of the 14 nucleotide substitutions observed in ten positions in post-whole-genome duplication yeast species. The library was transformed into *S. cerevisiae*, expressed under restrictive conditions and the enrichment of each genotype in the culture was quantified by deep sequencing before and after selection. After reprocessing of the raw data, we retained high quality fitness estimates for 3,847 variants (74.2%, see Methods).

Although the findings in [6] were based on the application of background-averaged epistasis theory, the prior limitation of the Walsh-Hadamard transform to only two alleles per sequence position required the authors to adopt an *ad hoc* strategy that involved performing separate analyses on combinatorially complete biallelic sub-landscapes. After imputing fitness values for missing variants we decomposed phenotypes into background-averaged epistatic coefficients using equation (9), which yielded very similar values for significant coefficients of all orders (S2b Fig, see Methods). However – as shown above for simulated data – our theory extensions permit the modeling of background-averaged epistasis in the context of multiallelic landscapes with missing data. We followed the same strategy as described in the previous section, using the tRNA DMS data to train Lasso regression models to predict variant fitness from sequence incorporating epistatic coefficients up to eighth order (Fig 2b, middle, see Methods). The resulting models include many higher-order epistatic coefficients (S1c Fig, ‘Background-averaged models’) yet exhibit extreme sparsity, with the median number of non-zero coefficients of any order ranging from 19 to 60 i.e. approximately 1% of all possible coefficients of eighth order or less (Fig 2b, middle). Models trained on at least 10% of the training data tend to explain more than 50% of the total explainable variance (Fig 2b, middle) and model-inferred background-averaged epistatic terms correlate well with those reported previously [6] (Pearson’s *r* = 0.79, Fig 2c, middle).

To evaluate whether the inferred models report on biologically relevant features of the underlying genetic landscapes, we tested whether sparse model coefficients were more likely to comprise genetic interactions (or modulators thereof) involving known physically interacting positions in the wild-type tRNA secondary structure (Fig 2a, middle). Regardless of data sparsity, background-averaged model coefficients tend to be significantly enriched for physical interactions (Fig 2d, middle). On the other hand, in the case of even moderate sub-sampling of training data (16%), one-hot model coefficients show no such enrichment (Fig 2d, middle). The results of this enrichment analysis closely mirror those obtained using simulated data (Fig 2d, left), suggesting that they are likely to generalise to other empirical genotype-phenotype landscapes.

Finally, we repeated similar analyses using a second DMS dataset comprising fluorescence measurements for all combinations of single substitution mutations at 13 positions (2^13^ = 8, 192 genotypes) separating two variants of the *Entacmaea quadricolor* protein eqFP611 (Fig 2, right column; see Methods) [7]. Lasso models capture the majority of fitness variance even with high levels of data sparsity (Fig 2b, right) and model-inferred background-averaged coefficients correlate well with those obtained using the decomposition in equation (9) (Pearson’s *r* = 0.83, Fig 2c, right). We also find that direct pairwise physical interactions involving the fluorophore (L63, Fig 2a, right) are enriched in model-derived background-averaged epistatic terms (Fig 2d, right), a result that is robust to missing data and recapitulates our observations obtained using ground truth genetic interactions in simulated DMS data.

## Discussion

We have provided an extension to the most rigorous computational framework available for describing and modeling empirical genotype-phenotype mappings. Beyond the study of background-averaged epistasis with respect to mutations in the primary sequence, this also permits the inclusion of ‘epimutations’ (changes in the epigenetic state of DNA), amino acid post-translational modifications or even particular environmental/experimental conditions.

In the simplest application, background-averaged epistatic coefficients (genetic interaction terms) can be directly computed from phenotypic measurements via the decomposition in equation (9). However, estimating epistatic coefficients by regression – as in equation (12) – is a more natural choice in the presence of missing data, when data for multiple related phenotypes is available [23] and/or in the presence of global epistasis [24, 25]. Our mathematical results provide three alternative methods to compute the multi-state (multiallelic) extension of the inverse Walsh-Hadamard transform *A*_*n*_, one of which allows the direct extraction of specific elements or sub-matrices. In which situations might this capability be desirable?

First, constructing full *A*_*n*_ matrices – particularly by numerical inversion – is impractical for large genetic landscapes. Second, the result of the product 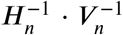 represents a matrix of sequence features when setting up the inference of epistatic (model) coefficients 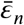 from phenotypic measurements 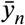as a regression task [23, 24, 26, 27]. The ability to construct this feature matrix in batches (of rows) allows computational resource-efficient iteration over large datasets when using frameworks such as TensorFlow or PyTorch.

Third, there are currently no methods to comprehensively map empirical genotype-phenotype landscapes with size greater than the low millions of genotypes. Therefore, assaying landscapes of this size or larger will typically involve experimental measurement of a (random) sub-sample of genotypes, corresponding to distinct rows in *A*_*n*_. In other words, it is usually unnecessary to construct full *A*_*n*_ matrices when modeling real experimental data. Finally, there is evidence of extreme sparsity in the epistatic architecture of biomolecules where only a small fraction of theoretically possible genetic interactions are non-zero [7]. The feasibility of sampling very large background-averaged epistatic coefficient spaces may improve methods to infer accurate genotype-phenotype models.

Using results from the analysis of both simulated and empirical multiallelic fitness landscapes, we have shown that sparse regression models relying on a background-averaged definition of epistasis can efficiently capture salient features of the underlying biological system – namely direct physical interactions or ground truth genetic interactions – even in situations of sparse sampling of phenotypes. This behaviour, which we speculate is due to a richer representation of the sequence feature space compared to one-hot models (i.e. higher level of constraint during model fitting; S1a Fig), is particularly desirable in the case of very large genetic landscapes where comprehensive phenotyping is infeasible. However, more work is needed to determine whether this result holds more generally. One difficulty in such comparisons between approaches is the requirement for a set of interactions or landscape features that are known to be critical for biomolecular function. Here we rely on Watson–Crick base pairing interactions and direct physical interactions involved in fluorophore orientation whose respective importance for RNA secondary structure and fluorescence activity are well-established.

More broadly, this work opens the door to investigations of the biological properties of background-averaged epistasis in empirical genetic landscapes of arbitrary shape and complexity. Beyond applications within the field of DMS, we believe our theory extensions have the potential to influence research in evolutionary and synthetic biology including protein engineering. In future it will be important to compare the performance and properties of models relying on this definition of epistasis to those of other recently proposed models that incorporate higher-order genetic interactions for phenotypic prediction [28, 29].

## Methods

### Simulated multiallelic fitness landscape

The simulated multiallelic genetic landscape (Fig 2a, left) consists of all possible combinations of 4 nucleotides per position in a 6-mer i.e. 4^6^ = 4, 096 possible genotypes. Consistent with observations of empirical fitness landscapes where single mutant phenotypes tend to be biased towards negative (detrimental) effects and larger in size than pairwise and higher-order epistatic terms, ground truth first-order epistatic coefficients (single nucleotide substitution effects) were drawn randomly from a normal distribution with a mean of -1 and standard deviation of 2 i.e. *ϵ* = *N*(*μ*, σ^2^) = *N*(−1, 4). Ground truth non-zero second-order terms (pairwise genetic interactions) were arbitrarily selected between the following pairs of positions: [1,2], [3,5] and [5,6], where their coefficient values were randomly sampled from *ϵ* = *N*(0, 1). All other second- and higher-order coefficients were set to zero. Fitness scores for all variants were calculated using equation (12). We simulated measurement error by adding Gaussian noise to all fitness scores of similar magnitude to the variance of first-order epistatic coefficients i.e. 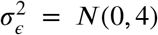. Finally, we also simulated versions of the same multiallelic fitness landscape with both relatively lower and higher measurement noise where 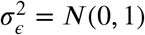 and 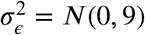, respectively (S1 Fig).

### Empirical combinatorial fitness landscapes

Raw sequencing (FASTQ) files obtained from the tRNA-Arg(CCU) DMS experiment in [6] were re-processed with DiMSum v1.3 [30] using default parameters with minor adjustments. We obtained fitness estimates for 5,059 out of a total of 5,184 possible variants (97.6%) in the combinatorially complete genetic landscape. We restricted the data to a high quality subset by requiring fitness estimates in all six biological replicates as well as at least 10 input read counts in all input samples. This resulted in a total of 3,847 retained variants (74.2%) for downstream analysis. For comparisons to previously-reported background-averaged coefficients (Fig 2c, S2b Fig), we used the author-processed data and analysis scripts [6]. Likewise, we used the author-processed data for imputation of missing phenotype values (replaced with the mean fitness at every mutation order) and calculation of background-averaged epistatic coefficients using the decomposition in equation (9) (S2b Fig). For the eqFP611 fluorescent protein DMS experiment, we used the author-processed fitness estimates (brightness scores) [7].

### Lasso regression models

We trained Lasso regression models to predict variant fitness estimates from nucleotide or amino acid sequences using the ‘scikit-learn’ Python package. Training data comprised random subsets of 1, 2, 4, 8, 16, 32 and 64% of retained variants of all mutation orders. All remaining held out variants comprised the ‘test’ data which was unseen during model training in each case.

To train models inferring background-averaged epistatic coefficients we used feature matrices of the form 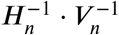(see equation (12)). For comparison, one-hot encoded matrices of sequence features were used. Linear regression was performed using 10-fold cross validation to determine the optimal value of the L1 regularization parameter λ in the range [0.005, 0.25] (‘LassoCV’ and ‘RepeatedKFold’ functions). Final models were fit to all training data. In order to estimate model-related statistics and performance results we fit 100 models to different random subsets of the training data for each model type and training data fraction. In Fig 2, S1 Fig and S2 Fig we plot the median of the indicated measures over all models, where error bars indicate the interquartile range. For model performance estimates in Fig 2b, the maximum explainable variance was calculated by substracting the total estimated technical variance (as reported by DiMSum fitness errors [30]) from the total fitness variance. For the simulated and eqFP611 DMS datasets, the maximum explainable variance was assumed to be 100%. In comparisons between sparse model-derived background-averaged epistatic coefficients and ground truth or previously-reported coefficients (scatter plots in Fig 2d and S1 Fig) we required model-derived terms to be non-zero in > 80% of models corresponding to random training data subsets.

To test enrichment of physical interactions in Lasso model coefficients we used the following approach: for each model, all position pairs represented in non-zero epistatic coefficients of at least second order were determined. The number of position pairs corresponding to direct physical interactions was counted and an associated enrichment score (odds ratio) and P-value calculated using Fisher’s Exact Test. The background set consisted of all position pairs in all possible epistatic coefficients. To test the appropriateness of the null hypothesis we also repeated enrichment analyses using random models i.e. randomly chosen sets of epistatic coefficients matching the numbers of non-zero coefficients in Lasso models and their distribution over different epistatic orders. The results of this analysis performed using the simulated dataset are shown in S1b Fig, left. Finally, the middle and right panels in S1b Fig assess the impact of measurement noise on the results of this enrichment analysis, revealing that noise level is anti-correlated with enrichment score and significance thereof, as expected.

## Acknowledgments

This work was funded by the Spanish Ministry of Science and Innovation (Severo Ochoa Centre of Excellence) and the CERCA Programme / Generalitat de Catalunya. A.J.F. was funded by a Ramón y Cajal fellowship (RYC2021-033375-I) financed by the Spanish Ministry of Science and Innovation (MCIN/AEI/ 10.13039/501100011033) and the European Union (NextGenerationEU/PRTR). B.L. was funded by European Research Council (ERC) Advanced grant (883742), the Spanish Ministry of Science and Innovation (LCF/PR/ HR21/52410004, EMBL Partnership), the Bettencourt Schueller Foundation, the AXA Research Fund and Agencia de Gestio d’Ajuts Universitaris i de Recerca (AGAUR, 2017 SGR 1322). V.M.P. was funded by the Spanish Ministry of Science and Innovation (PGC2018-100941-A-I00, PID2021-128976NB-I00). The funders had no role in study design, data collection and analysis, decision to publish, or preparation of the manuscript. We thank all members of the Lehner and Weghorn Labs for helpful discussions and suggestions.

## Supporting information captions

**S1 Fig. Supplementary figure related to Fig 2b**,**c**.

**S2 Fig. Supplementary figure related to Fig 2d**.

**S1 Text. Supplementary Methods**.

**Figure S1:**
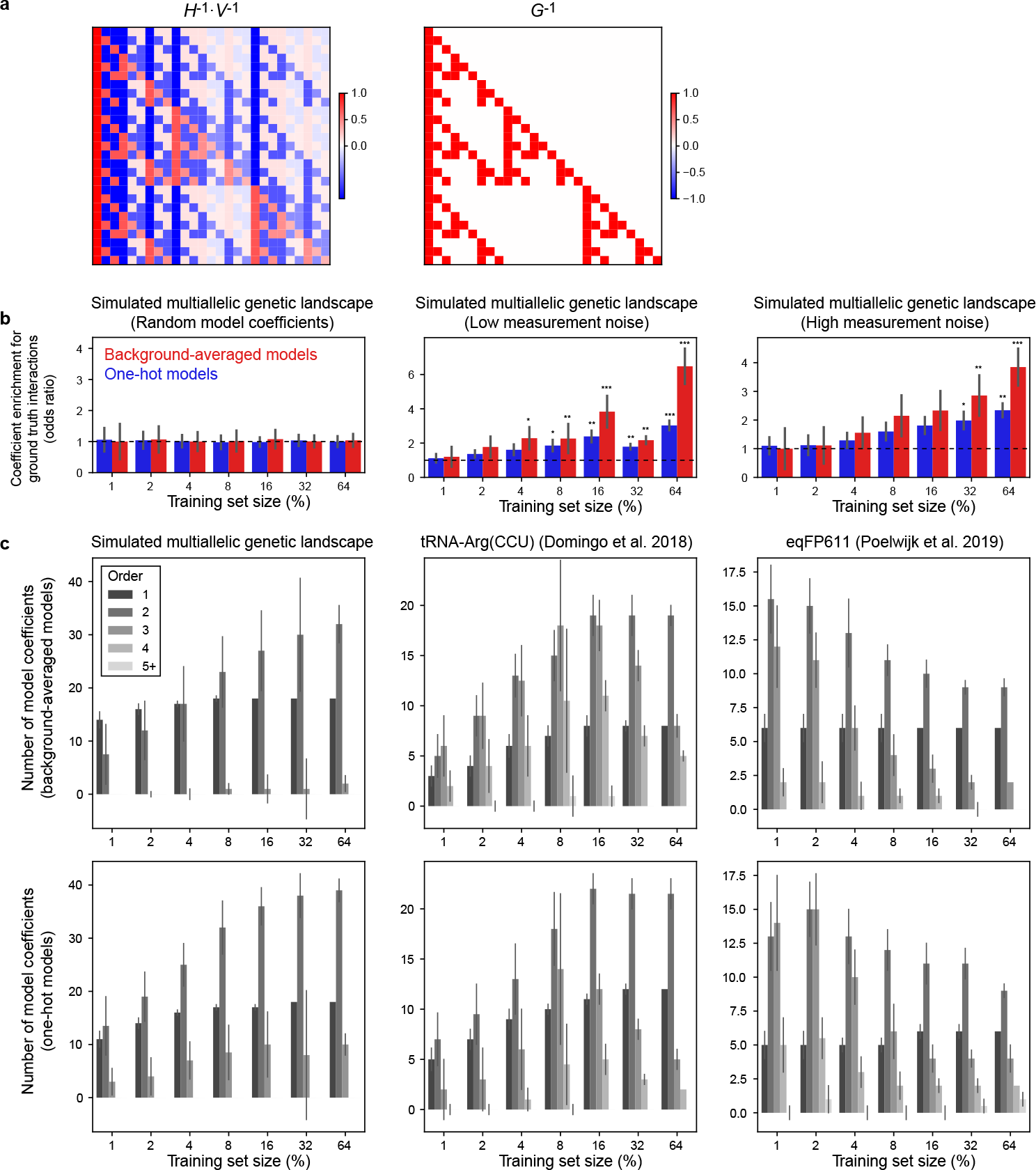
Supplementary figure related to Fig 2b,c. **a**, Cartoon depiction of alternative feature matrices for inferring epistatic coefficients by linear regression. *G*^−1^ in the right panel indicates the matrix of one-hot encoded sequence features – or embeddings – typically used when fitting models of genotype-phenotype landscapes [2]. The left panel represents the matrix of sequence features used to infer background-averaged epistatic coefficients, as in equation (12). **b**, Enrichment of direct physical interactions (see Fig 2a) in non-zero epistatic coefficients from sparse models of a simulated multiallelic fitness landscape. Left: results for random models with matching numbers of randomly selected epistatic coefficients at all coefficient orders (see Fig 2c, left). Middle, Right: results for sparse models fit to low and high measurement noise versions of the simulated fitness dataset, respectively. **c**, Numbers of non-zero espistatic coefficients of different orders in Lasso regression models inferred using different random fractions of the simulated and empirical combinatorial fitness landscapes indicated.

**Figure S2:**
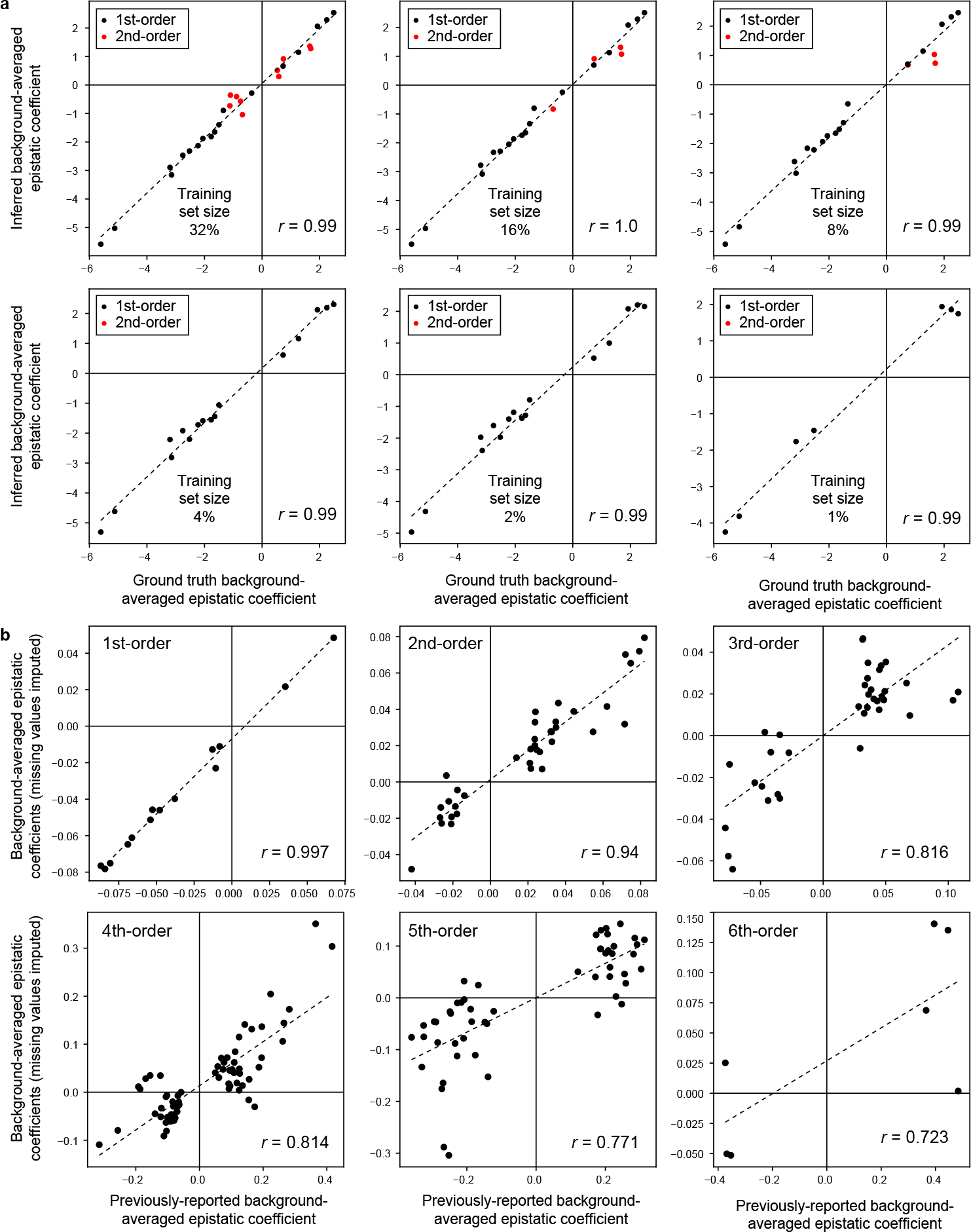
Supplementary figure related to Fig 2d. **a**, Scatter plots comparing sparse model-inferred background-averaged epistatic coefficients of a simulated multiallelic fitness landscape to ground truth values for varying training dataset sizes. **b**, Scatter plots comparing background-averaged epistatic coefficients obtained using the tRNA-Arg(CCU) DMS dataset and equation (9) to previously-reported significant coefficients [6] after imputing fitness values for missing variants (see Methods) and shown separately for coefficient orders 1-6. Only significant coefficients (FDR < 0.05) observed in at least 10 backgrounds are shown [6].

## Supporting Information 1 - Supplementary Methods

**Here we provide the proofs of the mathematical results shown in the main text**.

### Proposition 1.

*Let us define the matrices A*_*n*_ *recursively as*

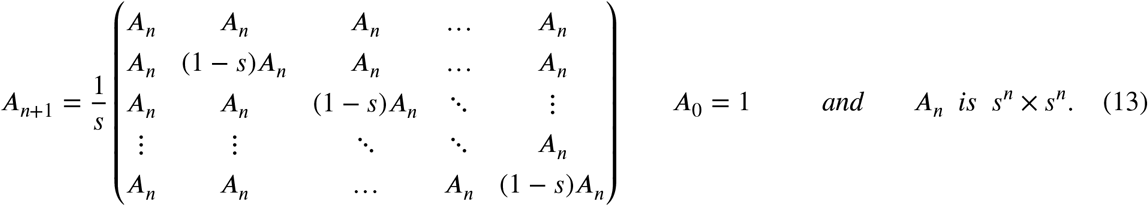

*For n* ∈ ℕ, *A*_*n*_ *is the inverse of the matrix H*_*n*_ *defined in equation (10)*.

*Proof*. Let us prove this by induction. For *n* = 1 we have

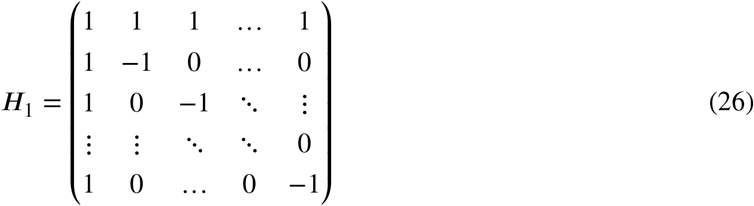

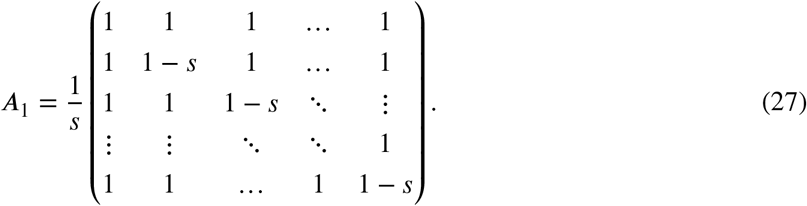

The rows and columns of these two matrices can be described as follows:

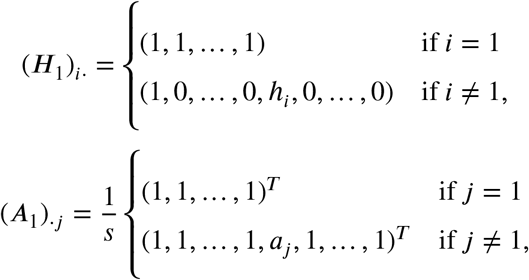

where *h*_*i*_ := (*H*_1_)_*ii*_ = −1 ∀*i* > 1 and a_*j*_ := (*A*_1_)_*jj*_ = 1 − *s* ∀*j* > 1.

Therefore,

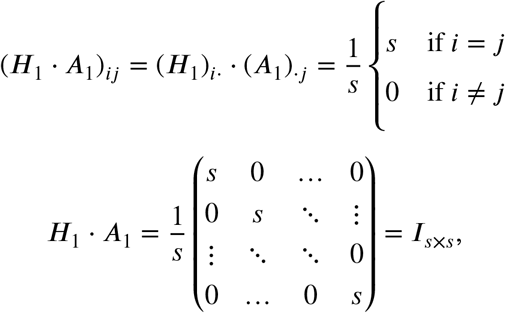

where *I*_*s*×*s*_ is the identity matrix of size *s* × *s*. Since both *H*_1_ and *A*_1_ are symmetric, it is also true that *A*_1_ ·*H*_1_ = *I*_*s*×*s*_. Therefore, *A*_1_ is the inverse of *H*_1_.

Assume that the hypothesis is true for a fixed *n* ∈ ℕ. Let us now prove that it is also true for *n* + 1. Following the recursive definitions of *H*_*n*+1_ and *A*_*n*+1_ in equations (10) and (13), we can write the blocks of these matrices as follows:

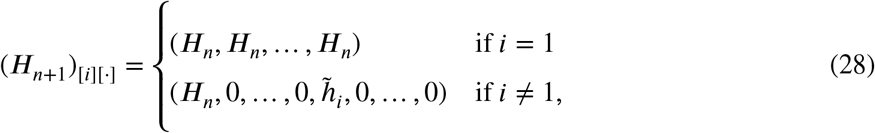

where (*H*_*n*+1_)_[*i*][*j*]_ denotes the block at position *i, j* in *H*_*n*+1_ and 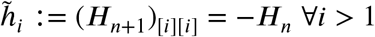;

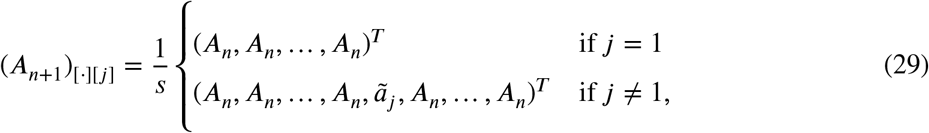

where (*A*_*n*+1_)_[*i*][*j*]_ denotes the block at position *i, j* in *A*_*n*+1_ and ã_*j*_ := *s*(*A*_*n*+1_)_[*j*][*j*]_ = (1 − *s*)*A*_*n*_ ∀*j* > 1. We can therefore write the block at position *i, j* of the product of these matrices as follows:

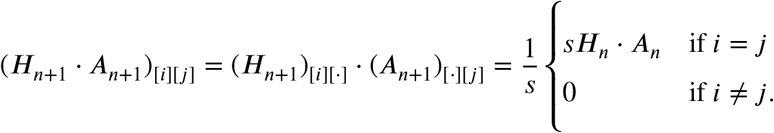

According to the induction hypothesis we know that *H*_*n*_ and *A*_*n*_ are inverse matrices i.e. 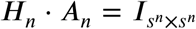. Therefore, the blocks on the diagonal are identity matrices and the blocks outside the diagonal are zeros. This means that 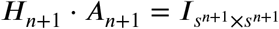. Similarly, due to the symmetry of the matrices we can also prove that *A*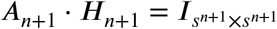.

We can then conclude that 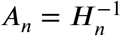 for every *n* ∈ ℕ.

### The proof above can be greatly simplified with the use of the Kronecker product and its properties as follows

*Proof*. It is straightforward to observe that the recursive notation of *H*_*n*+1_ can be expressed as a Kronecker product between *H*_1_ and *H*_*n*_. With this, we can use that the inverse of the Kronecker product is the product of the inverses of the individual matrices, if and only if, the individual matrices are invertible [31]. In our case, it is easy to see that the latter is true because we have shown that *H*_1_ is invertible (we computed its inverse in the previous proof), and all the other matrices are compute as a Kronecker product that derive uniquely from *H*_1_ (*H*_2_ is invertible because *H*_2_ = *H*_1_ ⊗ *H*_1_, *H*_3_ is invertible because *H*_3_ = *H*_1_ ⊗ *H*_2_, and so forth. Therefore,

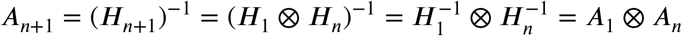

Now, we only need to compute the inverse of *H*_1_ as we did at the beginning of the original proof, and we obtain the recurrent definition of *A*_*n*+1_ (13) by applying the Kronecker product between *A*_1_ as in (27) and *A*_*n*_.

#### Proposition 2.

*The elements of H*_*n*_ *can be written as*

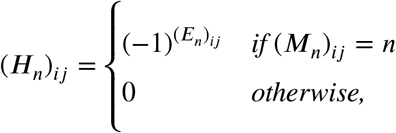

*where M and E are s*^*n*^ × *s*^*n*^ *matrices whose elements are*

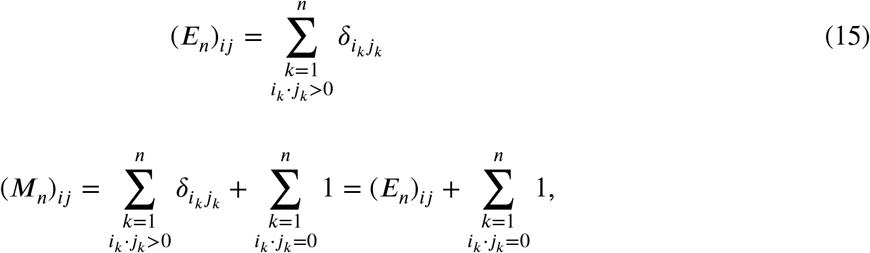

*where* δ_*ij*_ *denotes the Kronecker delta of i, j*.

*Proof*. Let us prove the formula by induction. For *n* = 1 and any given *s, H*_*n*_ is given by equation (26). Therefore, we can write (*E*_1_)_*ij*_ and (*M*_1_)_*ij*_ as follows:

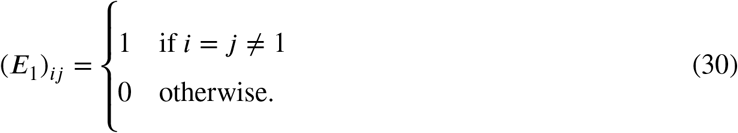

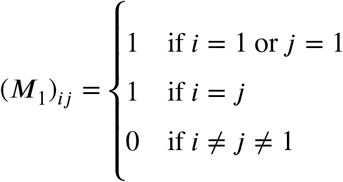

Therefore, since *n* = 1, (*M*_1_)_*ij*_ = *n* = 1 only when either *i* = 1, *j* = 1 or *i* = *j*. In the rest of the cases (*M*_1_)_*ij*_ ≠ *n* = 1 and, according to the formula, the elements of the matrix will be 0. Now, for the cases where (*M*_1_)_*ij*_ = *n* = 1, we need to check the value of (*E*_1_)_*ij*_. We can see how (*E*_1_)_*ij*_ = 1 only when *i* = *j* and they are different from 1. This means that all the elements of the diagonal of *H*_1_, except the first one, will be (−1)^1^ = −1 and the first row and first columns will have (−1)^0^ = 1. The rest of the elements correspond to (*M*_1_)_*ij*_ ≠ *n* = 1 so they will be filled with zeros. Putting all this together, we find the expression as *H*_1_ from equation (26).

Assume now that the expression is true for a fixed *n* ∈ ℕ and let us prove it for *n*+1. In this case, the matrix of *H*_*n*+1_ is defined by blocks (see equation (10)). We first define the indices *P* ∈ {1, …, *s*} and *Q* ∈ {1, …, *s*} for row and column blocks, respectively. The first matrix block of *H*_*n*+1_ corresponds to *P* = 1 and *Q* = 1. The corresponding blocks of the matrices *M*_*n*+1_ and *E*_*n*+1_, which are necessary for the derivation of the block in*H*_*n*+1_, are computed by comparing the genotype sequences 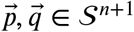 for which 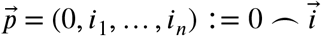 for 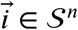 and 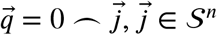. More generally, the block in the *P* ^*th*^ position with respect to the rows and *Q*^*th*^ position with respect to the columns can be obtained by comparing the genotype sequences 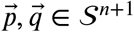 for which 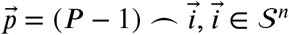 and 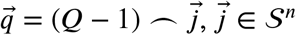. See below for a visual description of the notation.

From these observations, it can easily be deduced that for any 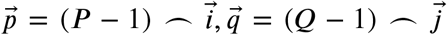 with 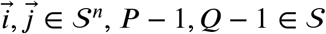,

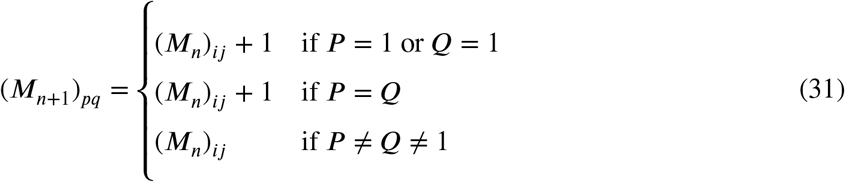

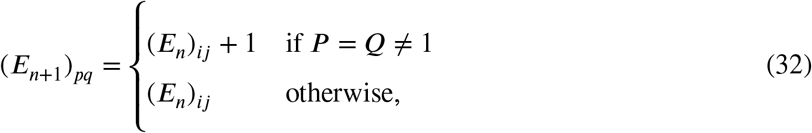

where *i* = *p* − *s*^*n*^(*P* − 1), *j* = *q* − *s*^*n*^(*Q* − 1), *P* = *p*/*s*^*n*^ and *Q* = *q*/*s*^*n*^, with. denoting the ceiling function. A visual description of the notation of the block structure of the matrices is given by

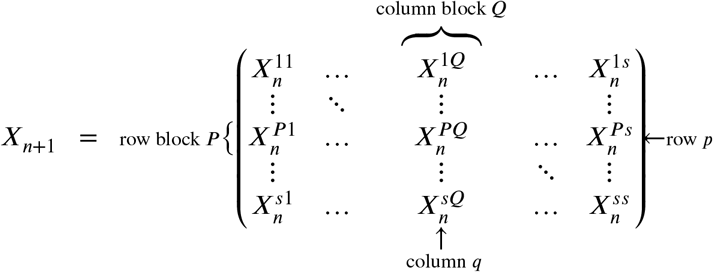

Here *X*_*n*+1_ denotes any generic matrix following the structure of the matrices in equations (10), (11), (13), (31) and (32).

Now, similar to the case *n* = 1, we have that (*M*_*n*+1_)_*pq*_ = *n* + 1 only when (*M*_*n*_)_*ij*_ = *n* and either *P* = 1, *Q* = 1 or *P* = *Q*, which corresponds to the newly added state being *p*_1_ = *P* − 1 = 0, *q*_1_ = *Q* − 1 = 0 or *p*_1_ = *q*_1_. In the rest of the cases (*M*_*n*+1_)_*pq*_ ≠ *n* + 1 and the elements of the matrix *H*_*n*+1_ will be 0. Now, for the cases where (*M*_*n*+1_)_*pq*_ = *n* + 1, we will have that (*E*_*n*+1_)_*pq*_ has either the same value of the corresponding entry in *E*_*n*_ or it will be increased by 1. This means, that when *P* = *Q* ≠ 1, the sign of the entry in *H*_*n*+1_ will be inverted. Otherwise, the sign of the element of the matrix stays the same. With this we prove the formula for *H*_*n*+1_, since we have shown how we can find the same block structure as in equation (10). By induction, we can conclude that the formula holds for every *n* ∈ ℕ.

#### Proposition 3.

*The elements of A*_*n*_ *can be written as*

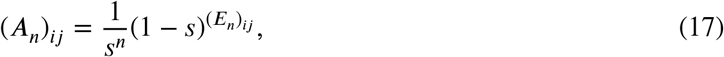

*where E*_*n*_ *is defined as in equation (15)*.

*Proof*. Let us prove the formula by induction.

For any given *s, A*_1_ is defined in equation 27. Its diagonal elements are equal to (1 − *s*)/*s* for *i* > 1, 1/*s* for *i* = 1 and its off-diagonal elements are equal to 1/*s*. It can easily be observed from equation (30) that the RHS of equation (17) is equal to (1 − *s*)/*s* for *i* = *j* ≠ 1 and to 1/*s* otherwise, so equation (17) is true for *n* = 1.

Assume now that equation (17) is true for a fixed *n* ∈ ℕ and let us prove it for *n* + 1. We use the recursive definition of *A*_*n*+1_ in equation (13) and the block representation of the genotype sequences as in the proof of (2).

Let us start with the first block in the diagonal of *A*_*n*+1_, i.e. *P* = 1 and *Q* = 1, where the entries of the corresponding block in *E*_*n*+1_ are derived from comparisons of pairs of genotype sequences of the form 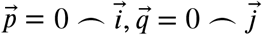 for 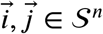. From equation (13), this block is equal to 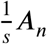, so writing *i* = *p* mod *s*^*n*^ and *j* = *q* mod *s*^*n*^, we have

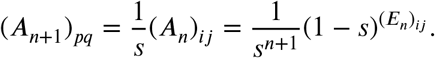

From equation (32), (*E*_*n*+1_)_*pq*_ = (*E*_*n*_)_*ij*_, which yields the desired result.

Now let us consider the elements in the other diagonal blocks of *A*_*n*+1_, where the entries correspond to pairs of genotype sequences of the form 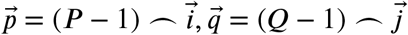 with *P* = *Q* ≠ 1 and 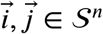. From equation (13), this block is equal to 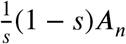, i.e.

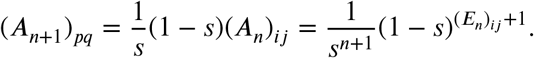

From equation (32), (*E*_*n*+1_)_*pq*_ = (*E*_*n*_)_*ij*_ + 1, which yields the desired result.

Finally, let us consider the elements in the off-diagonal blocks of *A*_*n*+1_, where the entries correspond to pairs of genotype sequences of the form 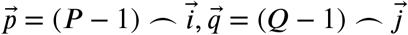 with *P* ≠ *Q* and 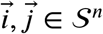. From equation (13), this block is equal to 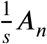, so we have

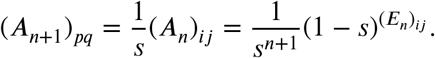

From equation (32), (*E*_*n*+1_)_*pq*_ = (*E*_*n*_)_*ij*_, which completes the proof.

#### Proposition 4.

*The matrices V*_*n*_ *and* 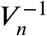*are diagonal matrices whose diagonal elements can be written as*

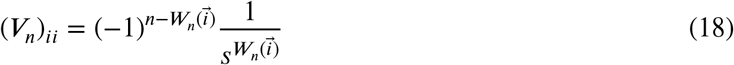

*and*

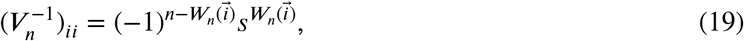

*where*

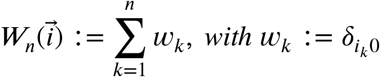

*and*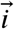 *again denotes the i*^*th*^ *element in 𝒮*^*n*^ *when ordered by the base s representation of integers*.

*Proof*. Let us prove equation (18) by induction, equation (19) follows directly.

One can easily check from equation (11) that the formula holds for *n* = 1. Let us now assume that equation (18) is true for a fixed *n* ∈ ℕ and let us prove it for *n* + 1. We use the recursive definition of *V*_*n*+1_ in equation (11). Let us consider the element (*V*_*n*+1_)_*pp*_. If 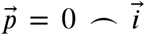, this corresponds to the first block of *V*_*n*+1_, i.e. *P* = 1, where the elements are multiplied by 1/*s* and 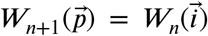, so

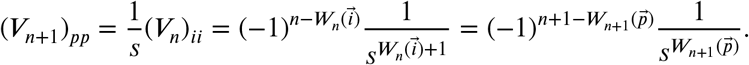

Similarly, if 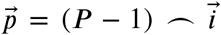 and *P* > 1, i.e. for the other diagonal blocks, from the recursive formula the elements are multiplied by −1 and 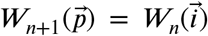, so by writing again *i* = *p* − *s*^*n*^(*P* − 1), where *P* = ⌈*p*/*s*^*n*^⌉, we have

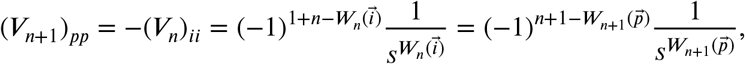

which completes the proof.

#### Proposition 5.

*The matrix A*_*n*_ *defined in equation (21) is the inverse of the matrix H*_*n*_ *in the general case where each position can have a different number of states*.

*Proof*. Let us prove by induction that *H*_*n*_ · *A*_*n*_ = *I* where *I* is the identity matrix of the corresponding size. Since *H*_*n*_ and *A*_*n*_ are symmetric, this would imply that *A*_*n*_ · *H*_*n*_ = *I* as well, and therefore, 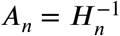.

The case *n* = 1 corresponds exactly to the case *n* = 1 of the proof of Proposition 1 by setting *s* = *s*_1_. Therefore, 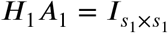, and *A*_1_ is the inverse of *H*_1_.

Now, assume the hypothesis is true for a fixed *n* ∈ ℕ and let us prove that this is also true for *n* + 1. We can write the rows and columns of the matrices *H*_*n*+1_ and *A*_*n*+1_ as equation (28) and equation (29), respectively. The only difference is that we need to replace *s* by *s*_*n*+1_ and the size of the matrices is different. Following exactly the same derivation as in Proposition 1 we can conclude that *H*_*n*+1_*A*_*n*+1_ = *I* and this proves by induction that 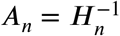.

#### Proposition 6.

*In this general case, the elements of H*_*n*_ *and A*_*n*_ *can be written as*

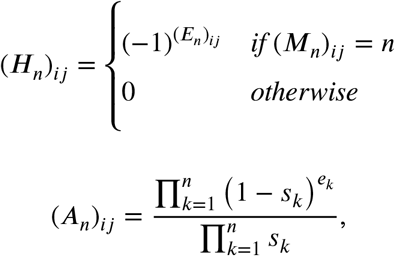

*where E*_*n*_ *and M*_*n*_ *are defined as in equation (15) and* 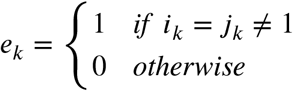

*Proof*. The proof follows directly from the proofs of Propositions 2 and 3. The only difference in the induction step is that *s* is replaced by *s*_*n*+1_.

#### Proposition 7.

*The matrices V*_*n*_ *and* 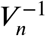 *are diagonal matrices whose diagonal elements can be written as*

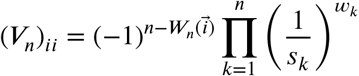

*and*

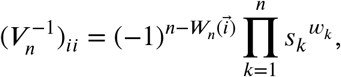

*where*

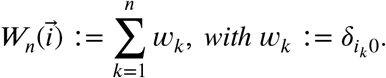

*Proof*. The proof follows the same steps as the proof of Proposition 4. The only difference in the induction step is that *s* is replaced by *s*_*n*+1_.

#### Proposition 8.

*Assuming that the uncertainty or empirical error of each component of* 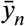 *is independent of the error of the other components, we can propagate the error to the estimation of the coefficients* 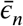 *as follows:*

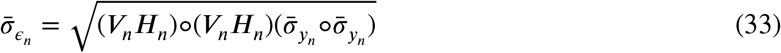

*where* 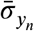 *and* 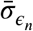 *denote the vector of uncertainties of the phenotypes and the epistatic coefficients, respectively, and* ° *denotes the element-wise product*.

*Proof*. According to equation (9) we can compute the epistatic coefficients as

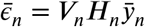

This means that each *ϵ* is a linear combination of the elements of 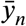, and the coefficients of the linear combination are defined by the elements of *V*_*n*_*H*_*n*_. Writing the formula as a system of equations and using that the error propagation of a linear function is just the square root of the sum of the squares of each coefficient times the variance of each variable, we obtain:

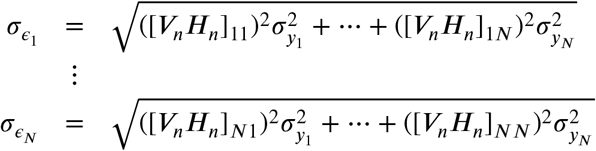

which can be written with matrix notation as

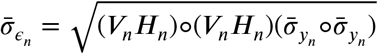

**This equation has already been used previously in articles regarding standard** *s* = 2 **epistasis [2], and it also applies here for any arbitrary** *n* **and** *s* **with our definition of** *V*_*n*_ **and** *H*_*n*_.

**For the example given in the main text regarding tRNA-ARG(CCU) with** *n* = 2 **and** *s* = 3, **we can see that applying the formula above we obtain:**

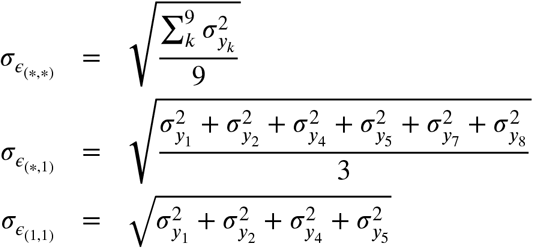

**where the other terms have the same shape as the corresponding same order term above, but with different phenotypes involved. We can see that if we assume equal uncertainties the error of the epistasis coefficients increase with the order of the interaction**.

